# NanoSimFormer: An end-to-end Transformer-based simulator for nanopore sequencing signal data

**DOI:** 10.64898/2026.01.20.700442

**Authors:** Shaohui Xie, Lulu Ding, Ling Liu, Zexuan Zhu

## Abstract

Nanopore sequencing has achieved a new standard of accuracy with the advent of R10.4.1 flow cell and high-performance Transformer-based basecalling models. However, existing signal simulators often fail to capture the complex, non-linear dynamics of nanopore current signals, relying on static pore models or lacking optimization objectives linked to basecalling, resulting in synthetic signals with substantially lower accuracy and fidelity than experimental data. To address this, we introduce NanoSimFormer, an end-to-end Transformer-based signal simulator that integrates basecaller guidance during training to generate high-fidelity nanopore signals explicitly optimized for accurate calling. Rigorous evaluation across diverse human, bacterial, and fungal R10.4.1 DNA sequencing datasets demonstrates that NanoSimFormer consistently outperformed competing methods (seq2squiggle and Squigulator), achieving median read accuracies exceeding 99% and Q-scores above 22.8, closely matching experimental baselines. NanoSimFormer faithfully recapitulated experimental variant calling performance on the human HG002 sample, achieving F1-scores of 0.9967 for SNPs and 0.8295 for small indels, and notably minimized false-positive errors in homopolymer and short tandem repeat (STR) regions where other simulators struggled. Furthermore, NanoSimFormer-derived reads enabled high-quality de novo bacterial assembly with consensus error rates below one mismatch per 100 kbp, comparable to experimental assemblies, and preserved fungal mock community structures with high correlation to experimental abundance profiles in metagenomic benchmarks. With tunable parameters for amplitude noise and event duration variance, NanoSimFormer enables the simulation of datasets spanning a wide range of data qualities. Together, these results establish NanoSimFormer as a robust tool for benchmarking and algorithm development in the latest nanopore sequencing era.

## Introduction

Nanopore sequencing has revolutionized genomics by enabling the direct, real-time analysis of long nucleic acid molecules without the need for amplification, a concept first established over three decades ago [1]. The technology’s unique ability to generate ultra-long reads has driven widespread adoption across diverse applications, such as single-nucleotide polymorphism (SNPs) and complex structural variants (SVs) detection [2, 3], de novo genome assembly [4], and modification calling [5]. Furthermore, the platform enables targeted analysis through “adaptive sampling” [6] and has recently demonstrated utility in clinical diagnostics for detecting disease-related biomarkers [7]. Recently, the field has witnessed a significant technological leap with the introduction of the R10.4.1 flow cell, characterized by a dual-reader head pore architecture, and the sequencing kit v14 utilizing the high-processivity E8.2.1 helicase enzyme [8]. When coupled with the latest official basecaller Dorado [9] and the released Transformer-based super-accuracy basecalling models (v5), this system achieves a new standard of performance, delivering modal raw read accuracies exceeding 99% (Q20+) [10].

As nanopore sequencing pushes the boundaries of accuracy and throughput, the demand for high-fidelity simulation tools has become critical. In silico data generation allows researchers to create infinite datasets under controlled scenarios for benchmarking analysis pipelines and enabling the debugging, optimization, and validation of novel algorithms in the absence of confounding experimental variables [11]. Historically, long-read simulators such as NanoSim [12], Trans-NanoSim [13], and Badread [14] have been instrumental in modeling error profiles for downstream tasks. However, these tools generate only nucleotide sequences and lack the raw ionic current signal, which is the essence of nanopore sequencing and has been proven to benefit the simulation performance greatly [15]. DeepSimulator [15, 16] pioneered this domain by leveraging a context-dependent model with a Bi-LSTM network, while NanosigSim [17] introduced Bi-directional Gated Recurrent Units to better capture signal noise. More recently, Squigulator [18] offered a computationally efficient approach using static k-mer pore models with empirical statistical distributions, and seq2squiggle [19] leveraged a non-autoregressive Feed-Forward Transformer to learn signal generation directly from training data.

Despite these advancements, existing nanopore signal simulation methods suffer from several critical limitations. First, these simulators are developed independently of the basecalling process and lack an optimization objective linked to basecalling. Without a feedback mechanism from the basecaller, these methods risk generating synthetic signals that are visually similar to real data but fail to retain the features essential for accurate decoding, thereby decoupling signal generation from the ultimate goal of high-fidelity basecalling. Second, most simulators typically rely on pre-existing k-mer pore models provided by Oxford Nanopore Technologies (ONT) to obtain the event level, such as DeepSimulator, NanosigSim, and Squigulator. Such reliance on static and discretized k-mer representations limits their capacity to model the complex, non-linear dynamics of nanopore current signals and prevents the establishment of long-range contextual relationships between signal and nucleotide sequences. Although simulators like Squigulator and seq2squiggle have introduced support for R10 chemistry, their modeling of the latest flow cell remains coarse and insufficiently optimized, yielding median read accuracies that are markedly lower than experimental data (91.91–95.29% as reported by the state-of-the-art seq2squiggle). To overcome the aforementioned issues, in this study, we propose NanoSimFormer, an end-to-end Transformer-based nanopore signal simulator that integrates basecaller guidance during training to generate high-fidelity signals tailored for the latest R10.4.1 chemistry. NanoSimFormer employs a non-autoregressive encoder-decoder architecture consisting of a Transformer-based sequence encoder and a hybrid Transformer-CNN signal decoder to capture complex contextual dependencies and non-linear signal dynamics. Unlike previous approaches, NanoSimFormer leverages a frozen basecalling model as a discriminator and is optimized via a multi-objective training strategy to produce signals specifically tuned for accurate basecalling. This end-to-end training pipeline derives sequence-to-signal mappings directly from the basecaller, thereby eliminating reliance on static k-mer pore models. NanoSimFormer was systematically evaluated against state-of-the-art simulators, seq2squiggle and Squigulator, across diverse datasets spanning human, bacterial, and fungal metagenomic samples. NanoSimFormer demonstrated superior signal fidelity, achieving median read accuracies (98.87% to 99.71%) and Q-scores (*>*22.8) that closely matched experimental baselines, whereas competing tools exhibited significant performance degradation. Importantly, the simulated reads supported high-precision downstream analysis, reproducing experimental performance in variant calling with F1-scores of 0.9967 for SNPs and 0.8295 for small indels, and facilitating de novo assemblies with consensus error rates below 1 mismatch per 100 kbp. Furthermore, NanoSimFormer provides tunable simulation parameters for amplitude noise and event duration variability, offering researchers flexible control to generate synthetic datasets with customizable quality profiles for diverse benchmarking scenarios.

## Results

### Overview of NanoSimFormer

NanoSimFormer utilizes an end-to-end Transformer-based architecture that integrates basecaller guidance to generate high-fidelity nanopore signals. During training (Fig. 1a), a frozen basecalling model processes experimental raw signals to derive the nucleotide sequences and low-resolution sequence-signal alignments (referred to as *moves* in the basecaller output), which map blocks of 6 signal timepoints to basecalled bases. To ensure consistency between training and inference, sequences are filtered using stringent criteria, including a minimum accuracy of 99% and a minimum quality score of Q20. The nucleotide sequences are mapped into high-level context features by a Transformer sequence encoder. These features are then temporally upsampled by duplicating each base-level feature multiple times, with the duplication factor determined by the extracted alignments. The resulting expanded features are subsequently passed to a hybrid Transformer-CNN signal decoder, which hierarchically increases the temporal resolution of the features to the scale of the raw signal. The model is trained using a multi-objective loss that combines a mean squared error (MSE) term for signal reconstruction and a basecalling loss calculated through a frozen basecalling model to ensure signal fidelity at both the signal and sequence levels. During simulation (Fig. 1b), where ground-truth alignments are unavailable, the upsampling process is driven by event durations randomly sampled from a duration sampler, and an amplitude noise sampler is applied to inject Gaussian noise into the decoded signal. NanoSimFormer supports diverse input formats, enabling the generation of synthetic signal files from either linear or circular genome references (FASTA), as well as experimental basecalled reads (FASTQ). More details on the implementation of the model architecture can be found in the Methods section and the Supplementary Materials. In the following sections, we demonstrate NanoSimFormer’s performance on real nanopore R10.4.1 datasets across various species, showing substantial improvements in basecalling accuracy and downstream analysis compared to existing tools.

**Figure 1.**
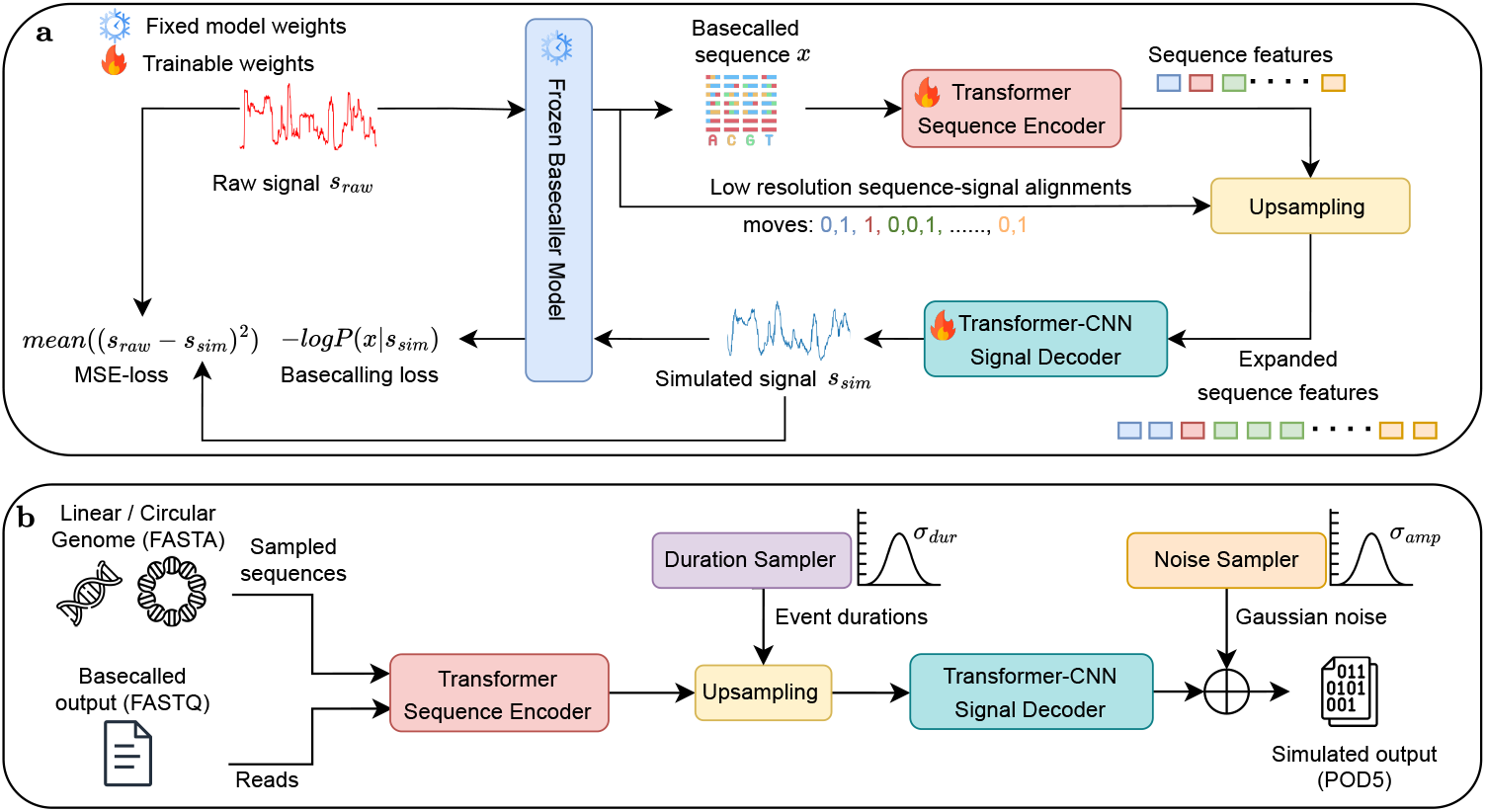
Overview of the NanoSimFormer framework. **a**, Training workflow incorporating base-caller guidance. The model utilizes a frozen basecaller (blue box with snowflake icon) to derive the basecalled sequence (*x*) and low-resolution sequence-signal alignments (“moves”, illustrated with different colors to indicate the signal duration associated with each base) from the raw signal (*s*_*raw*_). The trainable components (flame icon), comprising the Transformer-based sequence encoder (red) and Transformer-CNN signal decoder (teal). The high-level features from the sequence encoder are expanded via an upsampling operation, where features are duplicated according to the move information. Model is optimized using a multi-objective loss: mean squared error (MSE) loss that minimizes the discrepancy between the simulated signal (*s*_*sim*_) and *s*_*raw*_, and basecalling loss that minimizes the negative log-likelihood of the sequence given the simulated signal. **b**, Simulation workflow. For reference-based simulation, target sequences are randomly sampled from linear or circular genomes (FASTA). In contrast, for read-based simulation, basecalled reads (FASTQ) are utilized directly to generate a one-to-one simulated output. During simulation, a duration sampler generates event durations (with standard deviation *σ*_*dur*_) to replace the sequence–signal alignments used in training. Finally, a noise sampler adds Gaussian noise (with standard deviation *σ*_*amp*_) to the decoded signal to produce the final simulated output in POD5 format.

### Basecalling accuracy

To rigorously evaluate the signal fidelity of NanoSimFormer, a comprehensive benchmarking analysis was conducted against two state-of-the-art nanopore signal simulators, seq2squiggle (v0.3.4) [19] and Squigu-lator (v0.4.0) [18], which were selected as they represent leading approaches in the field and are among the few simulators that support the latest R10.4.1 pore chemistry used in our experiments. Experimental sequencing data were included in all comparisons to serve as the ground-truth baseline. Seq2squiggle was executed using its official pre-trained R10.4.1 model (trained on the reads of chromosomes 2, 3, and 4 of the human HCT116 dataset), while Squigulator was run using the default R10.4.1 DNA configuration. The NanoSimFormer model was trained on reads from chromosome 1 of a publicly available human HG002 R10.4.1 sample [20].

To assess generalization capabilities across diverse genomic contexts, all simulators were evaluated on a broad collection of independent R10.4.1 DNA sequencing datasets spanning human, bacterial, and fungal species, ensuring that none overlapped with the training data. For *Homo sapiens*, reads isolated from chromosome 22 of an HG002 R10.4.1 dataset were used for evaluation, while the corresponding simulation was performed using the perfectly accurate diploid chromosome 22 reference genome from the Telomere-to-Telomere consortium HG002 Q100 project (https://github.com/marbl/HG002). For bacterial evaluation, experimental datasets from five species were selected, including *Escherichia coli* (*E. coli*), *Klebsiella pneumoniae* (KP), *Morganella morganii* (MM), *Pseudomonas aeruginosa* (PA), and *Proteus mirabilis* (PM) [21]. Simulations for these bacterial species were conducted using high-quality ground truth reference genomes assembled with Hybracter (v0.11.2) using both ONT long reads and Illumina short-read data following the protocol described by [21]. Finally, for metagenomic evaluation, we used the ATCC Mycobiome Genomic DNA Mix (MSA-1010) dataset from the ont-open-data project [22], comprising a mixture of ten distinct fungal strains, including *Aspergillus fumigatus, Cryptococcus neoformans, Trichophyton interdigitale, Penicillium chrysogenum, Fusarium keratoplasticum, Candida albicans, Nakaseomyces glabratus, Malassezia globosa, Saccharomyces cerevisiae*, and *Cutaneotrichosporon dermatis*. Unlike the human and bacterial simulations described above, the simulations for MSA-1010 were conducted using experimental basecalled reads (FASTQ) as input, without reference-based read sampling, to preserve the natural concentration and relative abundance of the fungal community. To ensure a standardized comparison, simulated signals from all tools and the experimental raw signals were base-called using Dorado (v1.3.0) [9] with the super-accuracy model (dna r10.4.1 e8.2 400bps sup@v5.0.0). More details of the datasets and simulation settings can be found in the Methods section, as well as in Tables S1 and S2.

Basecalling performance was evaluated using multiple metrics, including read accuracy, PHRED quality score, mismatch rate, insertion rate, deletion rate, and read length (the length of a read excluding the left and right soft clips). As shown in Fig. 2 and Table S3, NanoSimFormer consistently generated signals with high fidelity, achieving median read accuracies ranging from 98.87% to 99.71%, closely matching the experimental range of 99.39% to 99.72%. In contrast, competing simulators showed substantial deviations from experimental data, with seq2squiggle and Squigulator yielding lower median accuracies, ranging from 93.34% to 97.25% and from 89.98% to 96.70%, respectively. This superiority in read accuracy is further evidenced by the PHRED quality scores. NanoSimFormer consistently achieved median Q-scores between 22.8 and 25.8 (indicating *>*99% base confidence), whereas seq2squiggle and Squigulator produced signals with markedly lower quality, consistently falling below a median Q-score of 16 across all datasets. NanoSimFormer maintained average median mismatch (0.21%), insertion (0.09%), and deletion (0.16%) rates that were remarkably close to the experimental baselines (0.19%, 0.06%, and 0.12%, respectively), whereas the other simulators frequently exhibited error rates exceeding 1.0% across these categories. NanoSimFormer also demonstrated robust generalization capabilities across both human and non-human datasets, whereas the performance of other simulators degraded substantially on non-human data. For instance, on the *Proteus mirabilis* (PM) dataset, NanoSimFormer achieved a median accuracy of 99.61%, closely matching the experimental accuracy of 99.65% and significantly outperforming seq2squiggle (94.72%) and Squigulator (91.91%). In the complex fungal metagenomic dataset (MSA-1010), all simulators exhibited reduced accuracy relative to reference-based simulations, as expected, because the simulation inputs were experimental basecalled reads that inherently contain sequencing errors that propagate into the generated signals. Despite this challenge, NanoSimFormer maintained a high median accuracy of 98.87%, remaining comparable to the experimental 99.40%. The observed accuracy difference (0.53%) is close to the intrinsic experimental read error (0.60%), demonstrating that NanoSimFormer faithfully preserves the sequence identity of the input reads. In contrast, seq2squiggle and Squigulator showed substantial performance degradation, with median accuracies of only 93.34% and 89.98%, respectively. While all simulators produced read length distributions broadly consistent with the experimental data, NanoSimFormer exhibited the closest overall agreement. Collectively, these results demonstrate the high fidelity, robustness, and generalization capability of NanoSimFormer for R10 chemistry signal simulation.

**Figure 2.**
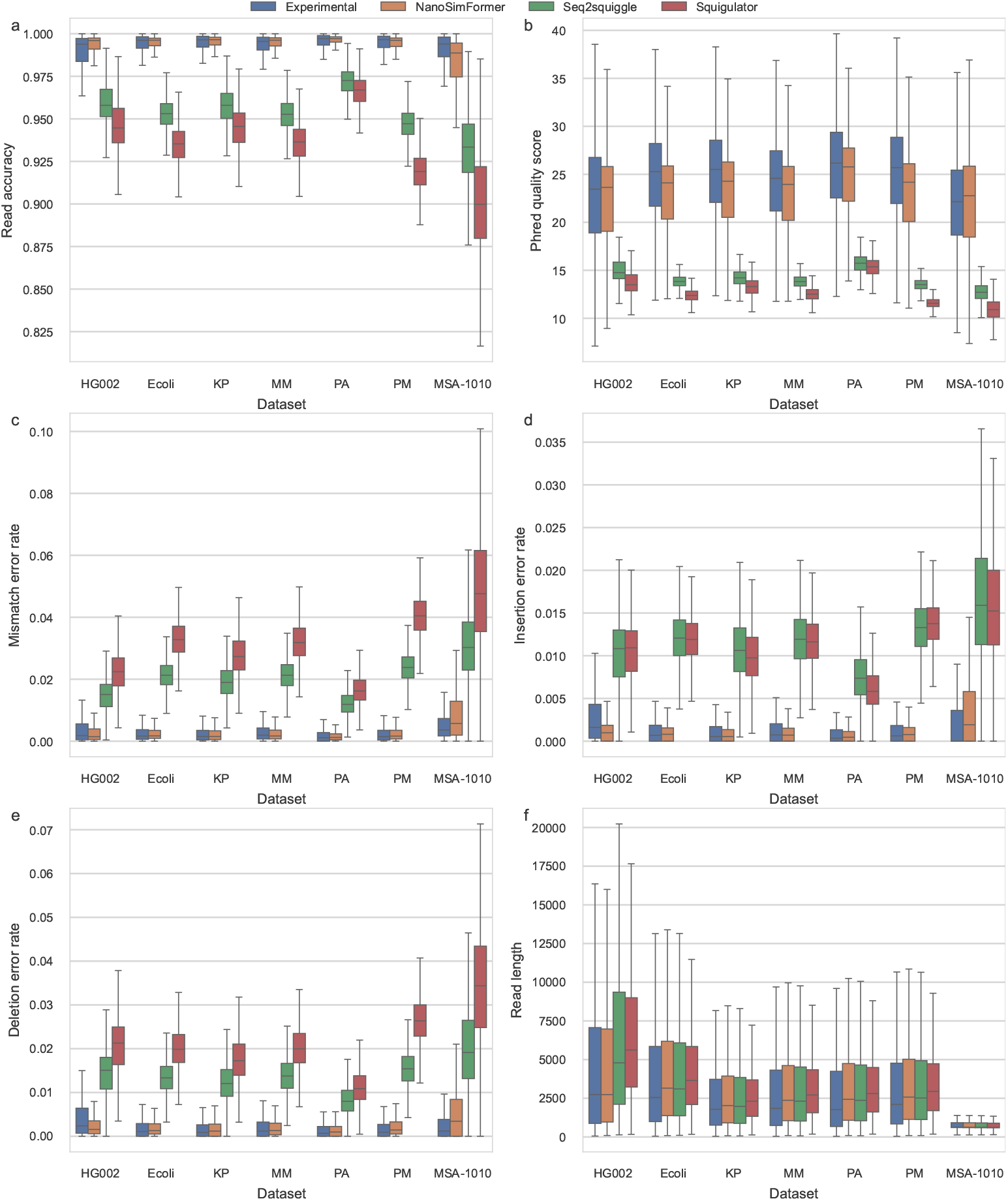
Basecalling performance comparison across various species. Performance metrics were evaluated for Experimental data (blue) and simulated signals from NanoSimFormer (orange), seq2squiggle (green), and Squigulator (red) across seven datasets. The evaluation metrics include: **a**, Read accuracy; **b**, Phred quality score; **c**, Mismatch error rate; **d**, Insertion error rate; **e**, Deletion error rate; and **f**, Read length. In the box plots, the center line represents the median, the edges of the box denote the first and third quartiles, thus enclosing the interquartile range (IQR), and the whiskers extend to the most extreme data points within 1.5 *×* IQR from the edges of the box. Sample reads used to derive these statistics are: HG002 (*n* = 320,000), E. coli (*n* = 254,210), KP (*n* = 365,393), MM (*n* = 261,589), PA (*n* = 211,667), PM (*n* = 140,727), and MSA-1010 (*n* = 1,884,844).

### Variant calling performance

To further assess the ability of simulators to faithfully recapitulate a complete nanopore sequencing analysis workflow, we rigorously evaluated variant detection performance following procedures established in previous studies [18, 19]. Both experimental and simulated reads derived from the HG002 diploid chromosome 22 reference were used for this evaluation. All reads were aligned to the GRCh38 reference genome using minimap2 (v2.28-r1209) [23]. Small variants, including single-nucleotide polymorphisms (SNPs) and small indels ≤30 bp, were called using Clair3 (v1.2.0) [24], while structural variants (SVs) were detected using Sniffles2 (v2.7.0) [25]. Small-variant call sets were benchmarked against the Genome in a Bottle (GIAB) [26] v3.3.2 high-confidence HG002 variants using RTGtools (v3.13) [27]. Structural-variant call sets were evaluated against the GIAB HG002 Structural Variants Draft Benchmark Set (draft benchmark V0.019-20241113 based on the T2T-HG002-Q100v1.1 diploid assembly) using Truvari (v5.4.0) [28].

NanoSimFormer demonstrated exceptional precision in SNP detection (Fig. 3a-b, Table S4), achieving an F1-score of 0.9967 (Precision: 0.9964, Recall: 0.9969), closely matching the performance of the experimental data (F1-score: 0.9979). In contrast, other simulators exhibited varying degrees of accuracy loss: seq2squiggle attained a moderate F1-score of 0.9657, whereas Squigulator showed a substantial performance decline with an F1-score of only 0.7486. Performance disparities were further pronounced in the detection of small indels, a challenging modality for nanopore sequencing. As detailed in Fig. 3c-d and Table S5, NanoSimFormer maintained a high indel F1-score of 0.8295, comparable to the experimental baseline of 0.8494. Conversely, seq2squiggle and Squigulator struggled to support accurate indel calling, yielding F1-scores of only 0.5060 and 0.2419, respectively. To elucidate the causes of the performance degradation observed in competing simulators, variant calling results were stratified by genomic context, with a particular focus on homopolymer and short tandem repeat (STR) regions. As shown in Fig. 4a, both seq2squiggle and Squigulator produced a high frequency of false-positive (FP) calls within homopolymer regions, whereas NanoSimFormer maintained a low FP rate comparable to that observed in experimental data. This finding is further supported by the homopolymer-specific error profiles presented in Figs. 4b-d, which quantify insertion, deletion, and mismatch rates across homopolymers of varying lengths in the HG002 dataset. NanoSimFormer consistently exhibited low error rates that closely tracked the experimental data across all homopolymer lengths. In contrast, seq2squiggle and Squigulator showed markedly elevated error rates—particularly deletion errors—that increased with homopolymer length (Fig. 4c). These substantial deviations from experimental error patterns in homopolymer regions likely lead to spurious indel calls during downstream analysis, thereby substantially limiting the suitability of seq2squiggle and Squigulator for robust variant-calling benchmarking. For structural variant detection, NanoSimFormer achieved an F1-score of 0.8437, comparable to the experimental data (0.8389) and outperforming both seq2squiggle (0.7918) and Squigulator (0.7970) (Table S6). Collectively, these findings demonstrate that NanoSimFormer is currently the sole simulator capable of generating synthetic signal datasets suitable for rigorous benchmarking in high-precision variant calling applications.

**Figure 3.**
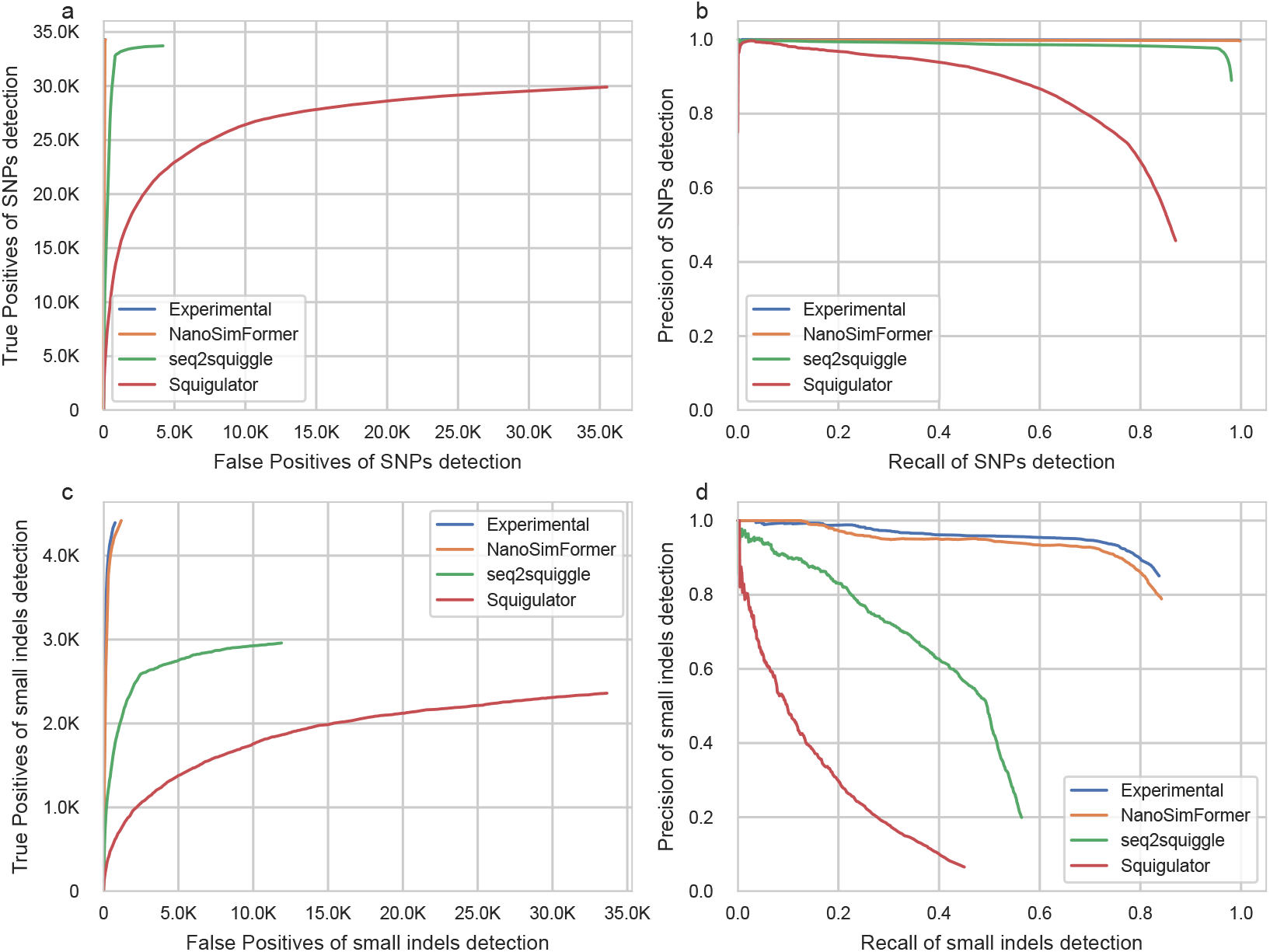
Small variant (SNPs and small indels) detection performance on the HG002 chromosome 22 dataset. **a**, Receiver Operating Characteristic (ROC) curves plotting the number of True Positives against False Positives for Single Nucleotide Polymorphisms (SNPs). **b**, Precision-Recall curves illustrating the trade-off between precision and recall for SNPs. **c**, ROC curves for small indels. **d**, Precision-Recall curves for small indels. Statistics for all panels were derived from the Human HG002 chromosome 22 dataset with a sample size of *n* = 320,000 reads.

**Figure 4.**
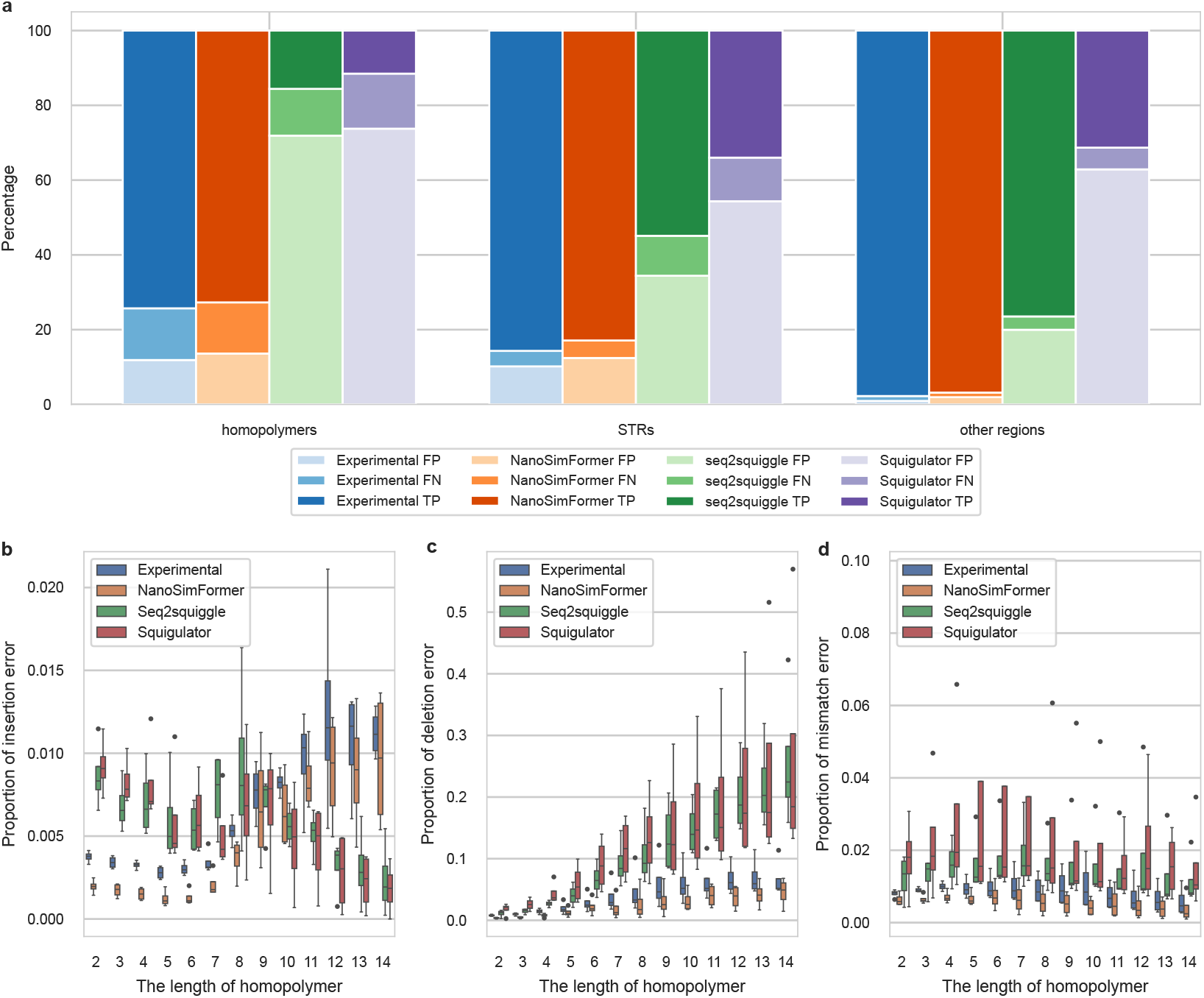
Percentage of detected small variants stratified by different genomic regions and the corresponding homopolymer error profiles on the HG002 chromosome 22 dataset. **a**, Stacked bar chart displaying the percentage of True Positives (TP, darkest shade), False Negatives (FN, medium shade), and False Positives (FP, lightest shade) of detected small variants stratified by genomic context, including homopolymers, short tandem repeats (STRs), and other genomic regions. **b-d** Insertion, deletion, and mismatch errors for homopolymers of varying lengths on the HG002 chromosome 22 dataset. Error proportions reflect the frequency of insertion, deletion, or mismatch errors occurring within these regions. For all box plots, the center line represents the median, the lower and upper bounds of the box represent the 25th and 75th percentiles, respectively, and the whiskers extend to the minimum and maximum values within 1.5 times the interquartile range, with outliers displayed as individual points. Statistics for all panels were derived from the Human HG002 chromosome 22 dataset with a sample size of *n* = 320,000 reads.

### Assembly performance

To assess the utility of the generated signals for downstream genomic assembly, de novo assemblies were performed on the five bacterial datasets (*E. coli, K. pneumoniae, M. morganii, P. aeruginosa*, and *P. mirabilis*). The simulated reads were assembled using nanoMDBG (v1.2) [29] and polished with Medaka (v2.1.1) [30]. The resulting contigs were evaluated against their corresponding reference genomes using QUAST (v5.2.0) [31]. As shown in Fig. 5a–c, NanoSimFormer produced assemblies with consensus quality comparable to that obtained from experimental data. At the base level, assemblies generated from NanoSimFormer and experimental reads consistently achieved near-perfect accuracy, with fewer than one mismatch or indel per 100 kbp across all evaluated datasets. In contrast, assemblies derived from seq2squiggle and Squigulator exhibited substantially higher error rates, with mismatch and indel densities ranging from tens to over a thousand per 100 kbp (e.g., Squigulator reached 1,067 mismatches per 100 kbp for *E. coli*). This discrepancy was further reflected in genome fraction coverage (Fig. 5c): NanoSimFormer assemblies achieved 99.90%–100.00% coverage, closely matching experimental results (99.99%–100.00%), whereas seq2squiggle and Squigulator frequently produced incomplete assemblies with lower and more variable coverage (e.g., 94.8% for Squigulator on *P. mirabilis*).

**Figure 5.**
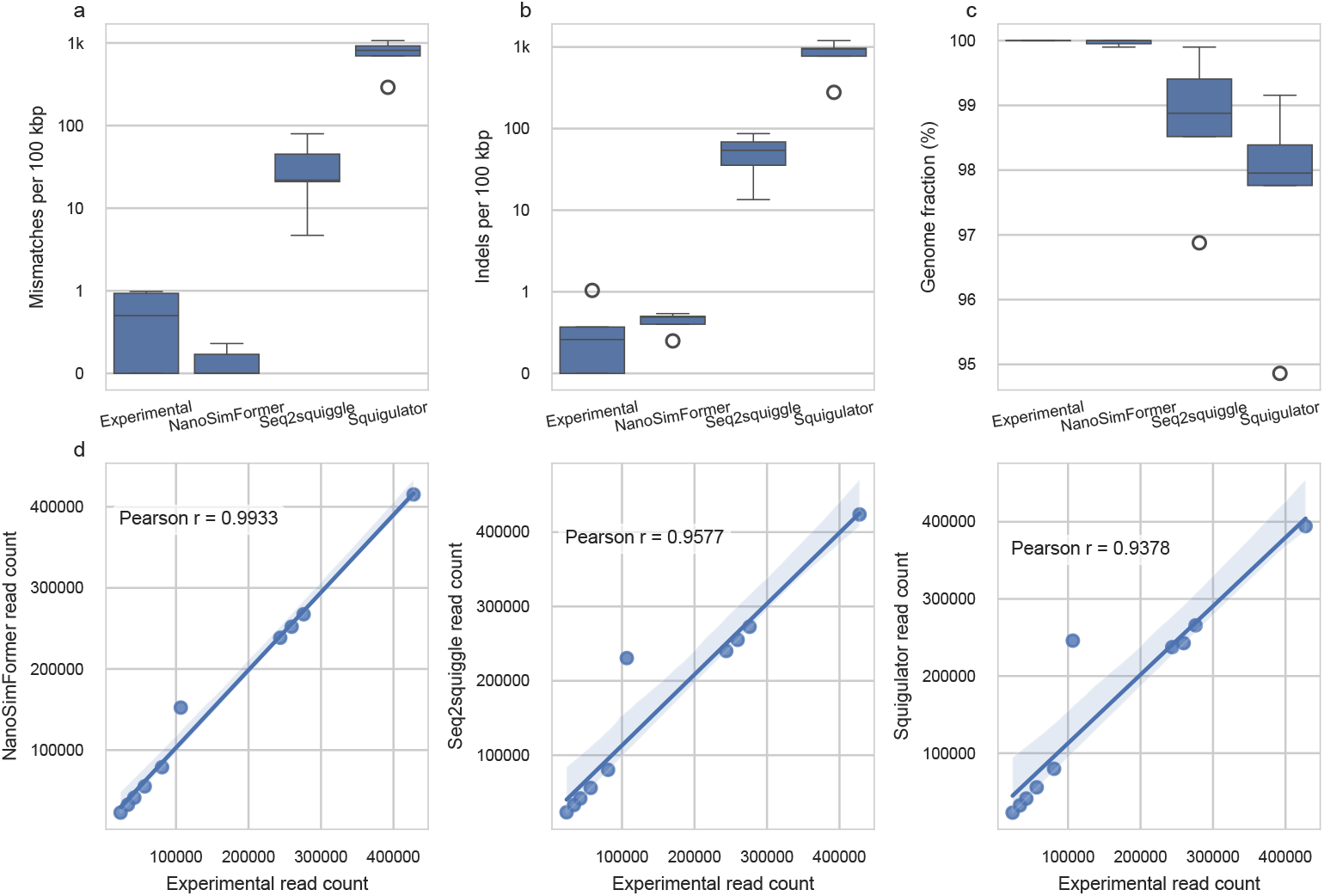
De novo assembly quality and metagenomic abundance preservation. De novo assembly performance was evaluated using simulated and experimental reads on *n* = 5 biologically independent bacterial datasets. The assemblies were assessed for **a**, Mismatches per 100 kbp; **b**, Indels per 100 kbp; and **c**, Genome fraction (%). For all box plots in panels **a, b** and **c**, the center line represents the median, the lower and upper bounds of the box represent the 25th and 75th percentiles, respectively, and the whiskers extend to the minimum and maximum values within 1.5 times the interquartile range, with outliers displayed as individual points. **d**, Scatter plots illustrating the correlation of read abundance (read counts classified by minimap2) between the experimental data and the simulated outputs for the MSA-1010 fungal mock community, comprising *n* = 10 distinct fungal strains. The solid blue line represents the linear regression fit, and the shaded region indicates the 95% confidence interval (CI) of the regression estimate. The Pearson correlation coefficient (*r*) is provided for each comparison.

For the MSA-1010 fungal metagenomic dataset, the preservation of community structure was evaluated by comparing the relative abundance of the ten constituent fungal strains between the experimental and simulated datasets. Read classification and taxonomic abundance estimation were performed by aligning basecalled reads to the MSA-1010 reference genome, which comprises the reference genomes of the ten known fungal strains in the sample, using minimap2. As illustrated in Fig. 5d, NanoSimFormer showed the strongest agreement with experimental data, achieving a Pearson correlation coefficient of 0.9933. This indicates that NanoSimFormer faithfully preserves the relative abundance and sequence identity of the input reads in complex metagenomic scenarios. In comparison, seq2squiggle (0.9577) and Squigulator (0.9378) showed lower correlations, suggesting greater deviation from the experimental profile during the simulation.

### Impact of tunable simulation parameters

To enable flexible generation of synthetic datasets with controllable accuracy for diverse benchmarking scenarios, we systematically analyzed the effects of two tunable simulation parameters: amplitude noise and event duration variance. The amplitude noise parameter adds zero-mean Gaussian noise with a tunable standard deviation, *σ*_*amp*_, to the simulated signal at the picoampere scale. The event duration parameter governs the distribution of sampled event durations, modeled as a Gaussian distribution with a fixed mean (*µ* = 2) and a tunable standard deviation (*σ*_*dur*_). These sampled durations correspond to low-resolution sequence–signal alignments, which map blocks of six signal timepoints to individual bases. Consequently, a mean duration of 2 corresponds to an average of approximately 12 signal samples per base, consistent with the R10.4.1 chemistry operated at 5 kHz sampling frequency and 400 bases per second.

As illustrated in Fig. 6, these parameters exhibited distinct effects on basecalling accuracy and downstream variant detection. For amplitude noise, NanoSimFormer demonstrated high stability within the standard deviation (*σ*_*amp*_) range of 0 to 1.5. As shown in Fig. 6a, read accuracy remained consistently high in this interval but exhibited a sharp decline as *σ*_*amp*_ exceeded 2.0. This trend was mirrored in variant detection performance. SNP detection maintained a high F1-score of *∼* 0.997 for *σ*_*amp*_ values ≤ 1.0, but dropped significantly to 0.9439 at *σ*_*amp*_ = 2.0 (Table S7, Fig. 6c, and Figs. S1a-b). A similar trend was also observed on the small indels detection (Table S8, Fig. 6d, and Figs. S1c-d).

**Figure 6.**
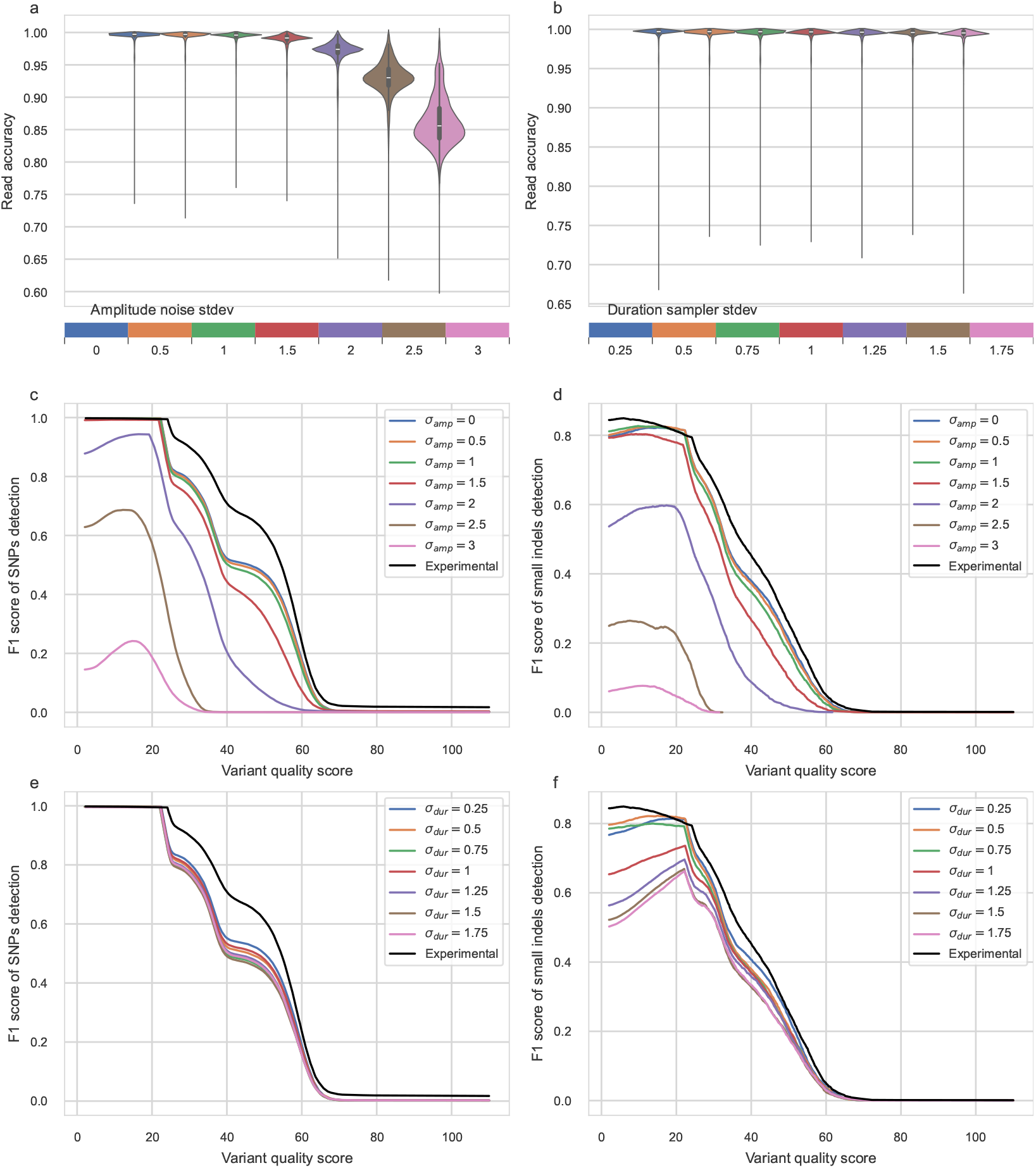
Impact of amplitude noise and event duration variance on signal fidelity and down-stream analysis. **a**,**b**, Violin plots illustrating the distribution of read accuracy for signals simulated by NanoSimFormer under varying parameters: **(a)** amplitude noise standard deviations (*Amp*.*σ*) ranging from 0 to 3, and **(b)** event duration standard deviations (*Dur*.*σ*) ranging from 0.25 to 1.75. In the violin plots, the internal box plots are defined as follows: the white center point represents the median, the lower and upper bounds of the thick bar represent the 25th and 75th percentiles, respectively, and the whiskers extend to the minimum and maximum values within 1.5 times the interquartile range. **c, d**, F1-score curves plotted against variant quality score thresholds for **(c)** SNPs and **(d)** small indels under varying amplitude noise intensities. **e, f**, F1-score curves plotted against variant quality score thresholds for **(e)** SNPs and **(f)** small indels under varying event duration variances. All statistics were derived from the Human HG002 chromosome 22 dataset with a sample size of *n* = 320,000 reads.

In contrast, variations in event duration exerted a more selective impact on performance metrics. Read accuracy remained remarkably robust across the entire tested range of duration standard deviations (from 0.25 to 1.75), showing minimal fluctuation (Fig. 6b). Similarly, SNP detection was largely insensitive to duration variance, maintaining F1-scores above 0.996 even at the highest tested standard deviation of 1.75 (Table S9, Fig. 6e and Figs. S2a-b). By contrast, indel detection sensitivity was inversely correlated with duration variance: the F1-score progressively decreased from 0.8227 (at *σ*_*dur*_ = 0.5) to 0.6626 (at *σ*_*dur*_ = 1.75) as the variance increased (Table S10, Fig. 6f and Figs. S2c-d). This sensitivity likely reflects the strong dependence of nanopore chemistry on dwell time for resolving homopolymer lengths. Smaller variance in dwell times—corresponding to more stable DNA translocation speeds—facilitates more accurate resolution for homopolymers and substantially lowers the false positives (FP) of small variants in homopolymer regions (Fig. S3). Based on these findings, a duration standard deviation of 0.5 was established as the default setting for the event duration sampler in NanoSimFormer, as it yielded the best overall balance in variant detection performance.

## Discussion

Nanopore sequencing has revolutionized genomics by enabling the analysis of long nucleic acid molecules, yet the development of accurate analysis algorithms relies heavily on high-fidelity simulation tools that can replicate the complex, non-linear dynamics of the latest flow cell chemistries. While recent advancements like the R10.4.1 flow cell have achieved raw read accuracies exceeding 99%, existing simulators often lag, relying on static pore models or decouple signal generation from the basecalling process. To address this gap, we developed NanoSimFormer, an end-to-end Transformer-based simulator that integrates basecaller guidance to generate signals explicitly optimized for accurate decoding. To assess the extent to which downstream analysis benefits from improved signal fidelity, we rigorously evaluated NanoSimFormer on diverse experimental datasets, including human, bacterial, and fungal samples, to prove that it significantly outperforms state-of-the-art simulators and faithfully recapitulates experimental performance across variant calling and assembly tasks.

Computational efficiency is a critical consideration for deep learning-based simulation tools. As detailed in Table S11, the computational cost of NanoSimFormer reflects the complexity of its Transformer-based encoder and hybrid Transformer-CNN decoder. Simulating the HG002 (chr22) dataset (320,000 reads, *∼* 40*×* depth) required 6,747 seconds with a peak memory usage of 5.67 GB (0.32 Mbp/s), a runtime comparable to the deep learning-based seq2squiggle (5,893 seconds) but significantly higher than the k-mer-based Squigulator (655 seconds, 3.26 Mbp/s). This performance highlights an inherent trade-off: while Squigulator offers rapid data generation suitable for high-throughput scenarios where coarse approximations suffice, the deep neural network approach of NanoSimFormer is necessary to capture the high-fidelity features required for precision benchmarking. Future architectural refinements, such as model quantization or knowledge distillation, could be applied to significantly reduce this computational overhead without compromising accuracy.

Looking forward, the flexible end-to-end architecture of NanoSimFormer provides a robust foundation for adaptation to emerging sequencing protocols and broader biotechnological applications. While the current model is optimized for R10.4.1 DNA chemistry, the basecaller-guided training strategy can be extended to Direct RNA Sequencing (DRS), such as the RNA004 protocol, to address the scarcity of high-fidelity transcriptomic simulators. Furthermore, because NanoSimFormer learns from raw signal data rather than fixed k-mer models, it is well-positioned for modification-based signal simulation. By integrating modification-aware basecallers (e.g., Dorado, *m*^6^*A*Basecaller [32]) into the training loop as discriminators, NanoSimFormer could be trained to generate signals that preserve patterns for specific epigenetic modifications like 5-methylcytosine (5mC) or N6-methylamine (6mA), thereby facilitating the development and benchmarking of modification detection algorithms. Beyond genomic benchmarking, this high signal fidelity opens avenues for specialized applications, such as the in silico design of sequencing barcodes to minimize de-multiplexing errors [33] and the rigorous stress-testing of error-correction codes for DNA data storage systems [34] against realistic signal noise profiles.

## Methods

NanoSimFormer employs an end-to-end Transformer-based encoder-decoder architecture designed to generate high-fidelity signals *s*_*sim*_ *∈* ℝ^*T*^ from an input nucleotide sequence *x ∈* Σ^*L*^, where *L* is the sequence length, *T* is the number of signal timepoints, and Σ = *{A, C, G, T}* is the alphabet of nucleotide bases.

### Sequence Encoder

The encoder of NanoSimFormer is designed to extract rich contextual representations from the nucleotide sequence (Fig. S4a). The input sequence *x* is first embedded into a high-dimensional space via an embedding layer. Specifically, each symbol is mapped to a vector using a learnable lookup table, resulting in a sequence of embeddings **h** *∈* ℝ^*L×d*^, where *d* is the feature dimension. These embeddings are then processed by *N* (*N* = 12 in our implementation) stacked Transformer encoder blocks. Each block comprises a Multi-Head Self-Attention (MHSA) module and a Gated Feed-Forward Network (FFN), both with layer normalization and residual connections. Given an input feature sequence **h**, the MHSA mechanism projects inputs into queries 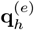, keys 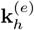, and values 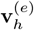 for each attention head *e ∈ {*1, …, *E}*, where *E* is the number of heads. Specifically:

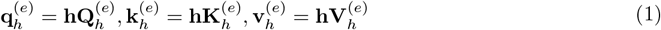

where 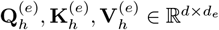 denote the learnable projection matrices that transform the signal feature sequence into queries, keys, and values, respectively. *d*_*e*_ is the head dimension (*d*_*e*_ = 64). Additionally, rotary positional embeddings (RoPE) [35] are then applied to the queries 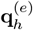 and keys 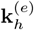 to incorporate relative positional information. Afterward, the self-attention weight 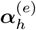 and the output representations 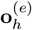 are computed for each head:

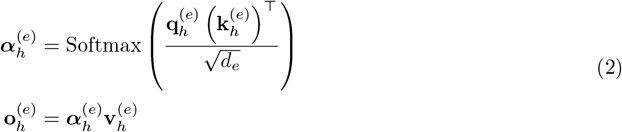

The outputs from all attention heads are concatenated and projected to the original dimension:

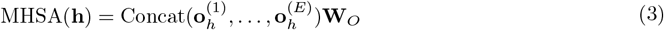

where **W**_*O*_ *∈* ℝ^(*E·de*)*×d*^ is a learnable output projection matrix. The FFN network after the MHSA module employs a Swish Gated Linear Unit (SwiGLU) variant [36]. The SwiGLU introduces nonlinearity, dimension expansion, and feature selection into the encoder layer. Specifically:

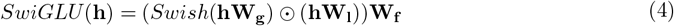

where **W**_**g**_, **W**_**l**_ *∈* ℝ^*d×*4*d*^ denote the learnable projection matrices that expand the dimension, whereas **W**_**f**_ *∈* ℝ^4*d×d*^ shrinks the dimension back to *d*.

### Upsampling process

Sequence features from the sequence encoder are upsampled by duplicating each base-level feature vector according to its corresponding event duration, analogous to the length regulator used in seq2squiggle [19]. During training, event durations are derived from the low-resolution sequence-signal alignments extracted from the basecaller output. In this alignment, one duration unit corresponds to a block of 6 signal timepoints (the stride of the basecaller). During simulation, event durations are instead generated by a duration sampler that models event lengths using a Gaussian distribution with a fixed mean (*µ* = 2, corresponding to an average of approximately 12 signal timepoints at the R10.4.1 5kHz sampling rate and 400bps translocation speed) and a tunable standard deviation *σ*_*dur*_ (default 0.5). This design effectively captures the stochastic variability of enzymatic motor stepping speed.

### Signal Decoder

The decoder of NanoSimFormer reconstructs the raw nanopore signal from the expanded sequence features using a hybrid Transformer-CNN architecture (Fig. S4b). The expanded features are first processed by *M* (*M* = 8 in our implementation) Transformer decoder blocks to capture signal-level dependencies. The resulting representations are then passed to a convolutional network that increases temporal resolution and regresses the scalar current values. This convolutional network consists of three sequential blocks, each comprising a transposed convolution layer, a ResNet block, and a Swish activation function. The channel dimensions progressively decrease from 256 to 128 and 64, while the transposed convolutional kernel sizes are 7, 5, and 6, with corresponding strides of 1, 2, and 3. Collectively, this design achieves a total 6-fold up-sampling of the temporal resolution relative to the input duration units. Finally, a standard convolution layer with a kernel size of 5 projects the 64-channel features to the final 1-dimensional continuous signal. During inference, an amplitude noise sampler adds zero-mean Gaussian noise with a tunable standard deviation (*σ*_*amp*_) to the decoded signal, thereby modeling the stochastic noise floor of the sensing hardware. For the default simulation setting, the standard deviation of amplitude noise is stochastically determined rather than fixed at a single value. Specifically, *σ*_*amp*_ is randomly sampled for each read from a Gamma distribution, which is designed to assign a lower noise level (typically between 0 and 1.0) to the majority of reads while subjecting a smaller proportion to higher noise levels (Fig S5). The parameters of this Gamma distribution were estimated using an Expectation-Maximization (EM) algorithm [37], which iteratively adjusted the noise standard deviation assignments to ensure that the resulting mixture of simulated read accuracies closely approximated the empirical accuracy distribution observed in experimental datasets.

### Optimization and loss functions

The model parameters *θ* are estimated by minimizing a multi-objective loss function *L*_*total*_ that enforces realism in both signal and sequence levels:

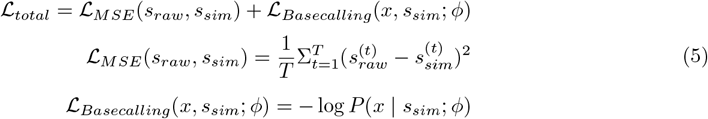

The mean squared error (MSE) loss ℒ_*MSE*_ measures the element-wise discrepancy between the simulated signal *s*_*sim*_ and the ground-truth raw signal *s*_*raw*_. The basecalling loss ℒ_*Basecalling*_ is defined as the negative log-likelihood of the nucleotide sequence *x* given the simulated signal. This is implemented using the connectionist temporal classification (CTC) loss [38] via a pre-trained and frozen basecaller (Dorado v5 super-accuracy model, parameterized by *ϕ*). During training, the basecaller parameters remain fixed, and gradients propagated through the basecaller affect only the sequence encoder and signal decoder.

### Experimental setup

#### Datasets

The NanoSimFormer model was trained on reads from chromosome 1 of a publicly available HG002 R10.4.1 sample (ENA accession: ERR12997168, from the gtgseq project) [20]. The sequencing reads were first basecalled using the Dorado super-accuracy model (dna r10.4.1 e8.2 400bps sup@v5.0.0) with the options `--emit-moves` to enable “moves” output, then aligned to the HG002 diploid reference genome (https://s3-us-west-2.amazonaws.com/human-pangenomics/T2T/HG002/assemblies/hg002v1.1.fasta.gz) to filter low-quality reads using stringent criteria, including a minimum accuracy of 99% and a minimum quality score of Q20. Subsequently, 13 million high-accuracy chunks with 5,000 signal samples were randomly selected from the dataset for training.

For simulation evaluation, we employed diverse samples and species strictly independent of the training set, including: *Homo sapiens* data from HG002 chromosome 22 reads [20], five species (*E. coli, K. pneumoniae, M. morganii, P. aeruginosa*, and *P. mirabilis*) with hybrid-assembled ground truth genomes [21], and the MSA-1010 fungal meta-genomic dataset comprising ten fungal strains [22].

#### Benchmark pipelines

In the simulation experiments, seq2squiggle and Squigulator were run using their default configuration. For reference-based simulation, NanoSimFormer selected an exponential distribution or a statistical model derived from the human HG002 sample as the read length distribution, based on an exploration of real experimental data. All simulators run with a mean read length configuration estimated from experimental data to ensure a similar sequencing depth as the experimental data. After basecalling, the reads were aligned to the respective reference genome using minimap2 (v2.28-r1209) [23] with the options “`-ax lr:hq --secondary=no --eqx`” to disable secondary alignments. The unaligned reads and the supplementary alignments were further excluded from the evaluation. The read accuracy is used as the primary metric for evaluating the basecalling performance:

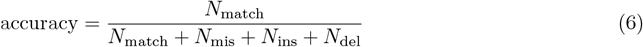

where *N*_match_, *N*_mis_, *N*_ins_, and *N*_del_, extracted from the CIGAR string, represent the numbers of matching bases, mismatches, insertions, and deletions of a read, respectively. The error (mismatch, insertion, and deletion) rate per read can be defined as:

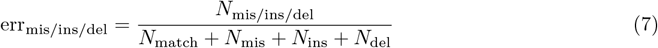

To rigorously assess the utility of simulated signals for downstream genomic analysis, we established comprehensive pipelines for variant detection and de novo assembly. For variant calling, basecalled reads were aligned to the GRCh38 reference genome using minimap2 (v2.28-r1209) [23]. Small variants (SNPs and indels) were identified using Clair3 (v1.2.0) [24] with the options `--include all ctgs` and `--platform=ont`. We employed the default Clair3 R10.4.1 super-accuracy model (`r1041 e82 400bps sup v500`) for experimental data, NanoSimFormer, and seq2squiggle. For Squigulator-simulated reads, we selected the Clair3 `r1041 e82 400bps sup v410` model as it achieved better results compared to the default selection. The resulting variant call files (VCFs) were filtered to exclude low-quality calls (QUAL*>*0 and DP*>*1) using bcftools and subsequently benchmarked against the GIAB v3.3.2 high-confidence variant set using RTG tools (v3.13) [27] vcfeval command. Structural variants (SVs) were detected using Sniffles2 (v2.7.0) [25] with tandem repeat annotations enabled. The SV call sets were evaluated against the GIAB HG002 Structural Variants Draft Benchmark using Truvari (v5.4.0) [28] with the parameters `--pick ac --passonly -r 2000 -C 5000`, followed by a refinement step (truvari refine) to optimize call coordinates. For de novo assembly, bacterial genomes were assembled using nanoMDBG (v1.2) [29] with the `--in-ont` option. The resulting contigs were polished using Medaka (v2.1.1) [30] and evaluated against ground-truth references using QUAST (v5.2.0) [31] to quantify metrics including genome fraction, indels, and mismatch rates.

#### Implementation and Environment configuration

All experiments were conducted on a Linux server equipped with two 48-core Intel® Xeon® PLATINUM 8558 CPUs (2.1 GHz), one NVIDIA H100 GPU, and 128 GB of system RAM. The operating system was Ubuntu 22.04.3 LTS. The implementation was developed using Python 3.10 and PyTorch 2.6.0 with CUDA 11.8 for GPU acceleration. NanoSimFormer was trained using gradient descent with the AdamW optimizer, an initial learning rate of 0.0002, and a batch size of 128. More details of the training configuration are listed in Table S12.

## Data availability

The sequencing data used in this study are available in the SRA/ENA databases and AWS Open Data registry under the accession codes and links listed in Supplementary Table S1.

## Code availability

The source code is available at GitHub: https://github.com/BioinfoSZU/NanoSimFormer.

## Acknowledgments

This study was supported by the National Key Research and Development Program of China under Grant 2022YFF1202104 and National Natural Science Foundation of China under Grant 62471310.

## Author Contributions

S.X. conceived and implemented NanoSimFormer. S.X. performed experiments, analyzed data, and drafted the manuscript. Z.Z. L.D. L.L. coordinated and supervised the project. L.D. L.L. and Z.Z. provided critical comments on algorithm evaluations and improved the manuscript.

## References

[1] Deamer, D., Akeson, M. & Branton, D. Three decades of nanopore sequencing. Nat. Biotechnol. 34, 518–524 (2016).

[2] Shafin, K. et al. Haplotype-aware variant calling with PEPPER-Margin-DeepVariant enables high accuracy in nanopore long-reads. Nat. Methods 18, 1322–1332 (2021).

[3] De Coster, W. et al. Structural variants identified by Oxford Nanopore PromethION sequencing of the human genome. Genome Res. 29, 1178–1187 (2019).

[4] Jain, M. et al. Nanopore sequencing and assembly of a human genome with ultra-long reads. Nat. Biotechnol. 36, 338–345 (2018).

[5] Simpson, J. T. et al. Detecting DNA cytosine methylation using nanopore sequencing. Nat. Methods 14, 407–410 (2017).

[6] Loose, M., Malla, S. & Stout, M. Real-time selective sequencing using nanopore technology. Nat. Methods 13, 751–754 (2016).

[7] Massaiu, I. et al. Accurate and rapid single nucleotide variation detection in PCSK9 gene using nanopore sequencing. Front. Med. 12, 1620405 (2025).

[8] Dorey, A. & Howorka, S. Nanopore DNA sequencing technologies and their applications towards single-molecule proteomics. Nat. Chem. 16, 314–334 (2024).

[9] Oxford Nanopore Technologies. Dorado. https://github.com/nanoporetech/dorado.

[10] Oxford Nanopore Technologies. Nanopore sequencing accuracy. https://nanoporetech.com/platform/accuracy (2026).

[11] Escalona, M., Rocha, S. & Posada, D. A comparison of tools for the simulation of genomic next-generation sequencing data. Nat. Rev. Genet. 17, 459–469 (2016).

[12] Yang, C., Chu, J., Warren, R. L. & Birol, I. NanoSim: nanopore sequence read simulator based on statistical characterization. GigaScience 6, gix010 (2017).

[13] Hafezqorani, S. et al. Trans-NanoSim characterizes and simulates nanopore RNA-sequencing data. GigaScience 9, giaa061 (2020).

[14] Wick, R. R. Badread: simulation of error-prone long reads. J. Open. Sour. Soft. 4, 1316 (2019).

[15] Li, Y. et al. DeepSimulator: a deep simulator for Nanopore sequencing. Bioinformatics 34, 2899– 2908 (2018).

[16] Li, Y. et al. DeepSimulator1. 5: a more powerful, quicker and lighter simulator for Nanopore sequencing. Bioinformatics 36, 2578–2580 (2020).

[17] Chen, W., Zhang, P., Song, L., Yang, J. & Han, C. Simulation of nanopore sequencing signals based on BiGRU. Sensors 20, 7244 (2020).

[18] Gamaarachchi, H., Ferguson, J. M., Samarakoon, H., Liyanage, K. & Deveson, I. W. Simulation of nanopore sequencing signal data with tunable parameters. Genome Res. 34, 778–783 (2024).

[19] Beslic, D. et al. End-to-end simulation of nanopore sequencing signals with feed-forward transformers. Bioinformatics 41, btae744 (2025).

[20] Wong, B., Ferguson, J. M., Do, J. Y., Gamaarachchi, H. & Deveson, I. W. Streamlining remote nanopore data access with slow5curl. GigaScience 13, giae016 (2024).

[21] Prior, K. et al. Accurate and reproducible whole-genome genotyping for bacterial genomic surveillance with Nanopore sequencing data. J. Clin. Microbiol. e00369–25 (2025).

[22] Oxford Nanopore Technologies. Oxford Nanopore Technologies Benchmark Datasets. https://registry.opendata.aws/ont-open-data (2024). Accessed: 2026-01-12.

[23] Li, H. Minimap2: pairwise alignment for nucleotide sequences. Bioinformatics 34, 3094–3100 (2018).

[24] Zheng, Z. et al. Symphonizing pileup and full-alignment for deep learning-based long-read variant calling. Nat. Comput. Sci. 2, 797–803 (2022).

[25] Smolka, M. et al. Detection of mosaic and population-level structural variants with Sniffles2. Nat. Biotechnol. 42, 1571–1580 (2024).

[26] Zook, J. M. et al. Extensive sequencing of seven human genomes to characterize benchmark reference materials. Sci. Data 3, 1–26 (2016).

[27] Cleary, J. G. et al. Comparing variant call files for performance benchmarking of next-generation sequencing variant calling pipelines. BioRxiv 023754 (2015).

[28] English, A. C., Menon, V. K., Gibbs, R. A., Metcalf, G. A. & Sedlazeck, F. J. Truvari: refined structural variant comparison preserves allelic diversity. Genome Biol. 23, 271 (2022).

[29] Benoit, G. et al. High-quality metagenome assembly from nanopore reads with nanoMDBG. BioRxiv 2025–04 (2025).

[30] Oxford Nanopore Technologies. Medaka. https://github.com/nanoporetech/medaka.

[31] Gurevich, A., Saveliev, V., Vyahhi, N. & Tesler, G. QUAST: quality assessment tool for genome assemblies. Bioinformatics 29, 1072–1075 (2013).

[32] Cruciani, S. et al. De novo basecalling of RNA modifications at single molecule and nucleotide resolution. Genome Biol. 26, 38 (2025).

[33] Li, J. et al. Empowering low-crosstalk, dynamic-decision random access of DNA storage via 384-multiplexed nanopore signatures. Nat. Commun. 16, 9233 (2025).

[34] Ding, L. et al. Polus: a Transformer-based Soft-decision Codec Enhancement Platform for DNA Storage. BioRxiv 2025–12 (2025).

[35] Su, J. et al. Roformer: Enhanced transformer with rotary position embedding. Neurocomputing 568, 127063 (2024).

[36] Shazeer, N. Glu variants improve transformer. arXiv preprint 2002.05202 (2020).

[37] Moon, T. K. The expectation-maximization algorithm. IEEE Signal Process. Mag. 13, 47–60 (1996).

[38] Graves, A., Fernández, S., Gomez, F. & Schmidhuber, J. Connectionist temporal classification: labelling unsegmented sequence data with recurrent neural networks. In Proc. 23rd Int. Conf. Mach. Learn., 369–376 (2006).

