## Supplementary Materials for "NanoSimFormer: An end-to-end Transformer-based simulator for nanopore sequencing signal data"

### Table S1. Evaluation dataset

| Dataset | Species | Flow cell | Library kit | Read number | Link or SRA/ENA accession ID | Link or NCBI accession ID of reference genome |
|---|---|---|---|---|---|---|
| HG002 (chr22) | *Homo sapiens* | FLO-PRO114M | SQK-LSK114 | 320000 | ERR12997168 | https://s3-us-west-2.amazonaws.com/human-pangenomics/T2T/HG002/assemblies/hg002v1.1.fasta.gz |
| E.coli | *Escherichia coli* | FLOMIN-114 | SQK-RBK114 | 254210 | ERR14711428 | https://zenodo.org/records/18220606/files/Ecoli_ref.fasta |
| KP | *Klebsiella pneumoniae* | FLOMIN-114 | SQK-RBK114 | 365393 | ERR14720565 | https://zenodo.org/records/18220606/files/Klebsiella_pneumoniae_ref.fasta |
| MM | *Morganella morganii* | FLOMIN-114 | SQK-RBK114 | 261589 | ERR14720737 | https://zenodo.org/records/18220606/files/Morganella_morganii_ref.fasta |
| PA | *Pseudomonas aeruginosa* | FLOMIN-114 | SQK-RBK114 | 211667 | ERR14720772 | https://zenodo.org/records/18220606/files/Pseudomonas_aeruginosa_ref.fasta |
| PM | *Proteus mirabilis* | FLOMIN-114 | SQK-RBK114 | 140727 | ERR14720750 | https://zenodo.org/records/18220606/files/Proteus_mirabilis_ref.fasta |
| MSA-1010 | Fungal mock community (10 fungal strains) | FLO-MIN114 | SQK-MAB114 | 1884844 | https://42basepairs.com/browse/s3/ont-open-data/fungal_ITS_2025.09/raw | ASM265v1, CNA3, ASM1935993v1, ASM71027v1, ASM2543354v1, ASM18296v3, CBS138_2MG, ASM18169v2, ASM4408963v1, ASM2001101v1 |

### Table S2. Simulator configuration

| Software Name (version) | Download url | Command |
|---|---|---|
| Squigulator (v0.4.0) | https://github.com/hasindu2008/squigulator | squigulator [reference] -x dna-r10-prom --seed 42 -r [mean_read_length] -o [output_blow5] -n [read_number] --ont-friendly=yes |
| Seq2squiggle (v0.3.4) | https://github.com/ZKI-PH-ImageAnalysis/seq2squiggle | seq2squiggle predict [reference] -o [output_blow5] --preserve-read-ids --num-reads [read_number] --read-length [mean_read_length] --seed 42 --profile dna-r10-prom --model R10_4_1.ckpt |
| NanoSimFormer (v1.0) | https://github.com/BioinfoSZU/NanoSimFormer | python nano_signal_simulator.py --input [reference] --output [output_pod5] --mode Reference --preset ont_r1041_dna_5khz --seed 42 --gpu [GPU_ID] --batch-size [batch_size] --sample-reads [read_number] |

### Table S3. Simulation basecalling performance

| **Dataset** | **Method** | **Read Accuracy (%)** | | **PHRED Quality Score** | | **Mismatch (%)** | | **Insertion(%)** | | **Deletion (%)** | | **Read Length (bp)** | |
|---|---|---|---|---|---|---|---|---|---|---|---|---|---|
| | | mean | median | mean | median | mean | median | mean | median | mean | median | mean | median |
| HG002 (chr22) | Experimental | 97.816 | 99.388 | 22.475 | 23.458 | 0.825 | 0.181 | 0.546 | 0.161 | 0.813 | 0.238 | 6636 | 2603 |
| | NanoSimFormer | **98.678** | **99.598** | 22.145 | **23.636** | **0.681** | **0.143** | **0.238** | **0.098** | **0.403** | **0.155** | 6839 | 2738 |
| | Seq2squiggle | 96.038 | 95.795 | 15.344 | 14.755 | 1.518 | 1.508 | 1.013 | 1.082 | 1.431 | 1.507 | 6804 | 4785 |
| | Squigulator | 94.773 | 94.467 | 14.02 | 13.491 | 2.239 | 2.243 | 1.020 | 1.093 | 1.969 | 2.128 | 6689 | 5617 |
| E.coli | Experimental | **99.043** | 99.617 | 24.84 | **25.272** | **0.398** | **0.163** | **0.227** | **0.068** | **0.331** | **0.107** | 4394 | 2525 |
| | NanoSimFormer | 98.734 | **99.622** | 22.495 | 24.108 | 0.660 | 0.171 | 0.243 | 0.081 | 0.363 | 0.129 | 4487 | 3153 |
| | Seq2squiggle | 95.263 | 95.303 | 13.871 | 13.835 | 2.152 | 2.129 | 1.223 | 1.207 | 1.362 | 1.333 | 4421 | 3099 |
| | Squigulator | 93.453 | 93.529 | 12.398 | 12.377 | 3.316 | 3.278 | 1.203 | 1.190 | 2.027 | 1.987 | 4347 | 3648 |
| KP | Experimental | **99.192** | 99.641 | 25.32 | **25.513** | **0.344** | **0.152** | **0.186** | **0.054** | **0.279** | **0.090** | 2767 | 1777 |
| | NanoSimFormer | 98.853 | **99.653** | 22.861 | 24.285 | 0.607 | 0.152 | 0.211 | 0.054 | 0.329 | 0.119 | 2863 | 2025 |
| | Seq2squiggle | 95.722 | 95.797 | 14.283 | 14.218 | 1.936 | 1.895 | 1.092 | 1.062 | 1.250 | 1.206 | 2801 | 1972 |
| | Squigulator | 94.402 | 94.558 | 13.271 | 13.299 | 2.808 | 2.732 | 1.008 | 0.975 | 1.782 | 1.725 | 2750 | 2308 |
| MM | Experimental | **99.013** | 99.552 | 24.314 | **24.583** | **0.410** | 0.195 | 0.234 | 0.073 | **0.342** | **0.120** | 3259 | 1833 |
| | NanoSimFormer | 98.758 | **99.622** | 22.447 | 23.952 | 0.647 | **0.168** | **0.229** | **0.071** | 0.366 | 0.131 | 3357 | 2366 |
| | Seq2squiggle | 95.232 | 95.266 | 13.876 | 13.826 | 2.152 | 2.129 | 1.211 | 1.193 | 1.405 | 1.376 | 3291 | 2311 |
| | Squigulator | 93.550 | 93.643 | 12.52 | 12.519 | 3.232 | 3.183 | 1.181 | 1.163 | 2.037 | 1.998 | 3233 | 2712 |
| PA | Experimental | **99.313** | **99.715** | 25.971 | **26.176** | **0.284** | **0.111** | **0.159** | **0.035** | **0.244** | **0.068** | 3356 | 1760 |
| | NanoSimFormer | 99.060 | 99.711 | 24.382 | 25.776 | 0.492 | 0.121 | 0.185 | 0.046 | 0.263 | 0.097 | 3446 | 2430 |
| | Seq2squiggle | 97.130 | 97.251 | 15.746 | 15.737 | 1.247 | 1.189 | 0.775 | 0.737 | 0.849 | 0.798 | 3380 | 2367 |
| | Squigulator | 96.520 | 96.700 | 15.293 | 15.363 | 1.706 | 1.619 | 0.624 | 0.583 | 1.150 | 1.088 | 3342 | 2804 |
| PM | Experimental | **99.089** | **99.652** | 25.351 | **25.7** | **0.380** | **0.143** | **0.220** | **0.062** | **0.312** | **0.092** | 3557 | 2078 |
| | NanoSimFormer | 98.662 | 99.614 | 22.554 | 24.177 | 0.685 | 0.163 | 0.242 | 0.077 | 0.411 | 0.144 | 3646 | 2575 |
| | Seq2squiggle | 94.692 | 94.715 | 13.543 | 13.503 | 2.398 | 2.377 | 1.345 | 1.329 | 1.564 | 1.539 | 3587 | 2515 |
| | Squigulator | 91.893 | 91.908 | 11.603 | 11.571 | 4.065 | 4.046 | 1.384 | 1.376 | 2.658 | 2.637 | 3514 | 2944 |
| MSA-1010 (read mode) | Experimental | **98.499** | **99.395** | 21.9 | 22.141 | **0.724** | **0.360** | **0.322** | **0.000** | **0.456** | **0.120** | 604 | 534 |
| | NanoSimFormer | 97.516 | 98.868 | 21.726 | **22.774** | 1.245 | 0.571 | 0.501 | 0.192 | 0.738 | 0.344 | 607 | 535 |
| | Seq2squiggle | 93.017 | 93.341 | 12.764 | 12.708 | 3.203 | 3.025 | 1.694 | 1.591 | 2.086 | 1.913 | 614 | 535 |
| | Squigulator | 90.016 | 89.981 | 10.988 | 10.903 | 4.902 | 4.762 | 1.595 | 1.524 | 3.487 | 3.436 | 608 | 528 |

**Table S4. HG002 chr22 SNPs detection performance (under the threshold with the best F1 score) of different simulators**

| **method** | **variant quality score threshold** | **f1** | **precision** | **recall** | **precision-recall AUC** |
|---|---|---|---|---|---|
| Experimental | 3.84 | **0.9979** | **0.9977** | **0.9980** | 0.9886 |
| NanoSimFormer | 7.11 | 0.9967 | 0.9964 | 0.9969 | **0.9919** |
| seq2squiggle | 17.07 | 0.9657 | 0.9740 | 0.9574 | 0.9685 |
| Squigulator | 15.32 | 0.7486 | 0.7536 | 0.7436 | 0.7669 |

**Table S5. HG002 chr22 small indels detection performance (under the threshold with the best F1 score) of different simulators**

| method | variant quality score threshold | f1 | precision | recall | precision-recall AUC |
|---|---|---|---|---|---|
| Experimental | 5.78 | **0.8494** | **0.8801** | **0.8207** | **0.8055** |
| NanoSimFormer | 10.93 | 0.8295 | 0.8653 | 0.7965 | 0.7969 |
| seq2squiggle | 19.35 | 0.5060 | 0.5482 | 0.4699 | 0.4037 |
| Squigulator | 18.27 | 0.2419 | 0.2549 | 0.2302 | 0.1458 |

### Table S6. Performance of structural variant detection on HG002 chr22 dataset

| Methods | TP | FP | FN | Precision | Recall | F1 |
|---|---|---|---|---|---|---|
| Experimental | 384 | 26 | 136 | 0.9366 | 0.7597 | 0.8389 |
| NanoSimFormer | 391 | 26 | 132 | 0.9376 | 0.7668 | 0.8437 |
| Seq2squiggle | 371 | 54 | 156 | 0.8729 | 0.7244 | 0.7918 |
| Squigulator | 362 | 29 | 170 | 0.9258 | 0.6996 | 0.7970 |

**Table S7. HG002 chr22 SNPs detection performance (under the threshold with the best F1 score) for NanoSimFormer simulated signal using different amplitude noise variance.**

| amplitude noise stdev | variant quality score threshold | f1 | precision | recall | precision-recall AUC |
|---|---|---|---|---|---|
| 0 | 9.17 | **0.9977** | 0.9978 | 0.9975 | **0.9959** |
| 0.5 | 10.28 | 0.9977 | 0.9980 | 0.9973 | 0.9958 |
| 1 | 9.53 | 0.9974 | 0.9977 | 0.9970 | 0.9951 |
| 1.5 | 14.59 | 0.9933 | 0.9937 | 0.9928 | 0.9923 |
| 2 | 17.63 | 0.9439 | 0.9403 | 0.9474 | 0.9502 |
| 2.5 | 12.26 | 0.6869 | 0.6676 | 0.7073 | 0.6803 |
| 3 | 15.57 | 0.2424 | 0.2568 | 0.2296 | 0.1240 |

**Table S8. HG002 chr22 small indels detection performance (under the threshold with the best F1 score) for NanoSimFormer simulated signal using different amplitude noise variance.**

| amplitude noise stdev | variant quality score threshold | f1 | precision | recall | precision-recall AUC |
|---|---|---|---|---|---|
| 0 | 15.17 | 0.8227 | 0.8844 | 0.7691 | 0.7814 |
| 0.5 | 13.16 | 0.8264 | 0.8730 | 0.7845 | 0.7860 |
| 1 | 10.07 | **0.8271** | 0.8515 | 0.8040 | **0.7891** |
| 1.5 | 9.35 | 0.8038 | 0.8186 | 0.7895 | 0.7680 |
| 2 | 17.55 | 0.5977 | 0.5824 | 0.6138 | 0.5387 |
| 2.5 | 7.56 | 0.2648 | 0.1918 | 0.4277 | 0.1511 |
| 3 | 11.16 | 0.0769 | 0.0557 | 0.1241 | 0.0111 |

**Table S9. HG002 chr22 SNPs detection performance (under the threshold with the best F1 score) for NanoSimFormer simulated signal using different event duration variance.**

| event duration stdev | variant quality score threshold | f1 | precision | recall | precision-recall AUC |
|---|---|---|---|---|---|
| 0.25 | 11.13 | 0.9975 | 0.9975 | 0.9974 | 0.9954 |
| 0.5 | 9.17 | **0.9977** | 0.9978 | 0.9975 | **0.9959** |
| 0.75 | 8.75 | 0.9973 | 0.9973 | 0.9972 | 0.9954 |
| 1 | 8.47 | 0.9970 | 0.9971 | 0.9969 | 0.9951 |
| 1.25 | 5.94 | 0.9965 | 0.9965 | 0.9964 | 0.9941 |
| 1.5 | 10.20 | 0.9965 | 0.9972 | 0.9957 | 0.9937 |
| 1.75 | 6.82 | 0.9960 | 0.9963 | 0.9956 | 0.9929 |

**Table S10. HG002 chr22 small indels detection performance (under the threshold with the best F1 score) for NanoSimFormer simulated signal using different event duration variance.**

| event duration stdev | variant quality score threshold | f1 | precision | recall | precision-recall AUC |
|---|---|---|---|---|---|
| 0.25 | 19.86 | 0.8148 | 0.9045 | 0.7412 | 0.7664 |
| 0.5 | 15.17 | **0.8227** | 0.8844 | 0.7691 | **0.7814** |
| 0.75 | 13.68 | 0.8004 | 0.8455 | 0.7599 | 0.7624 |
| 1 | 22.30 | 0.7360 | 0.8100 | 0.6744 | 0.6673 |
| 1.25 | 22.19 | 0.6961 | 0.7391 | 0.6578 | 0.6073 |
| 1.5 | 22.09 | 0.6689 | 0.6974 | 0.6425 | 0.5673 |
| 1.75 | 22.22 | 0.6626 | 0.6893 | 0.6379 | 0.5574 |

**Table S11. Runtime and memory usage.**

| Dataset | Simulators | Time (s) | Speed (Mbp/s) | Memory usage (GB) |
|---|---|---|---|---|
| HG002 (chr22) | NanoSimFormer | 6747 | 0.32 | 5.67 |
| | Seq2squiggle | 5893 | 0.37 | 5.89 |
| | Squigulator | 655.81 | 3.26 | 0.32 |
| E.coli | NanoSimFormer | 3753 | 0.30 | 4.70 |
| | Seq2squiggle | 3494.09 | 0.32 | 4.32 |
| | Squigulator | 405.49 | 2.73 | 0.20 |
| KP | NanoSimFormer | 3561.97 | 0.29 | 4.48 |
| | Seq2squiggle | 3354.74 | 0.30 | 3.95 |
| | Squigulator | 224.27 | 4.48 | 0.17 |
| MM | NanoSimFormer | 2901.18 | 0.30 | 4.41 |
| | Seq2squiggle | 2219.24 | 0.39 | 3.85 |
| | Squigulator | 189.36 | 4.47 | 0.16 |
| PA | NanoSimFormer | 2404.6 | 0.30 | 4.21 |
| | Seq2squiggle | 2043.85 | 0.35 | 3.64 |
| | Squigulator | 195.56 | 3.62 | 0.18 |
| PM | NanoSimFormer | 1902.24 | 0.27 | 4.05 |
| | Seq2squiggle | 1803.95 | 0.28 | 3.34 |
| | Squigulator | 108.28 | 4.57 | 0.17 |
| MSA-1010 | NanoSimFormer | 6626 | 0.23 | 5.79 |
| | Seq2squiggle | 4572 | 0.33 | 6.55 |
| | Squigulator | 525 | 2.83 | 2.21 |

### Table S12. Training setting

| | |
|---|---|
| Batch size | 128 |
| Signal chunk length | 5000 |
| Maximum nucleotide sequence length (bp) | 500 |
| Chunk number | 13403827 |
| Step size | 104717 |
| Feature dimension | 512 |
| Attention head number | 8 |
| Sequence Transformer encoder layers | 12 |
| Signal Transformer decoder layers | 8 |
| Optimizer | AdamW |
| Optimizer betas | (0.9, 0.999) |
| Optimizer eps | 1e-08 |
| Optimizer weight decay | 0.01 |
| Initial learning rate | 2e-4 |
| Learning rate scheduler | Linear Warmup Cosine Annealing |
| Warmup steps | 500 |
| Automatic mixed precision | Float16 |
| Gradient norm clip | 1.0 |

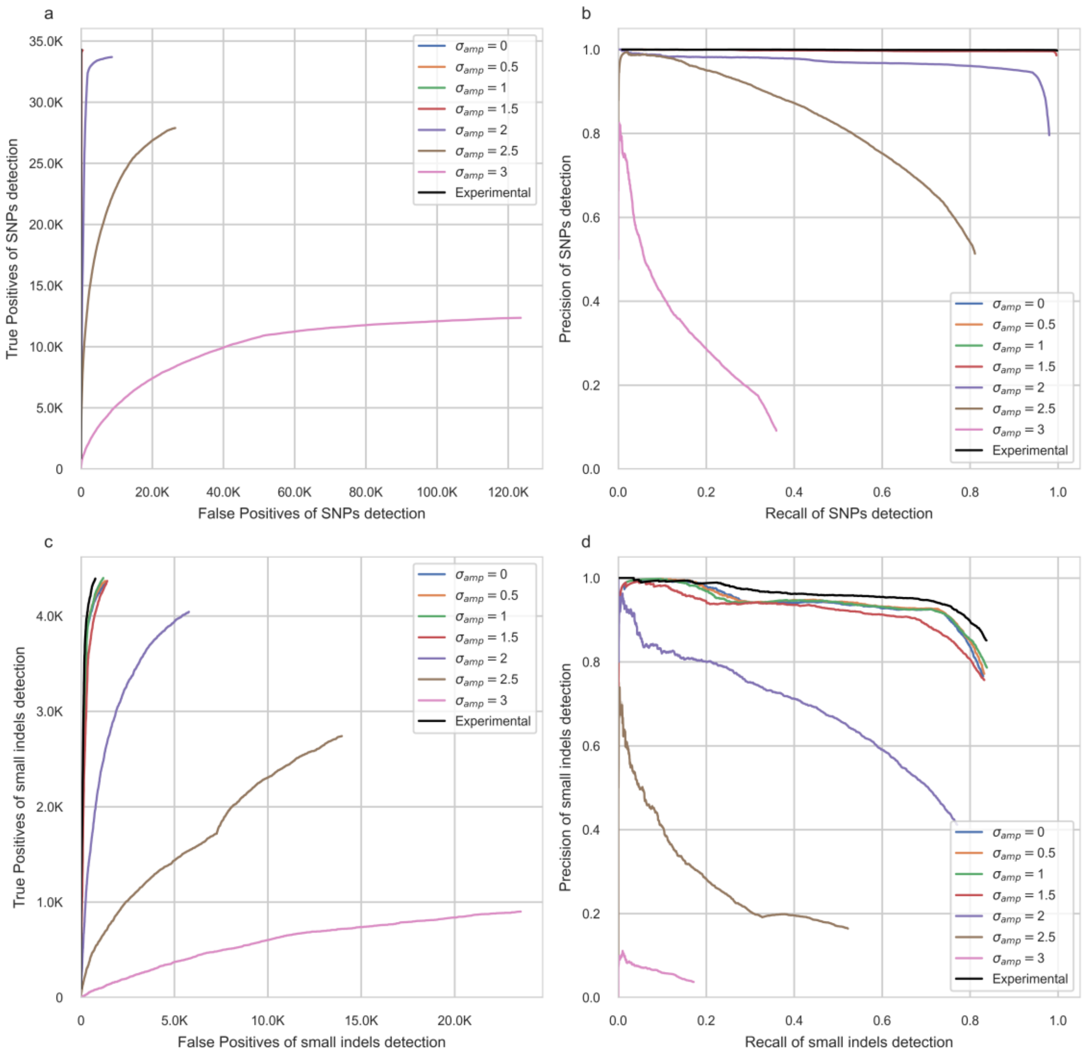

**Fig S1. Small variants detection performance for NanoSimFormer simulated signal using different amplitude noise variance. a,** ROC curves for SNPs. **b,** Precision-recall curves for SNPs. **c,** ROC curves for small indels. **d.** Precision-recall curves for small indels.

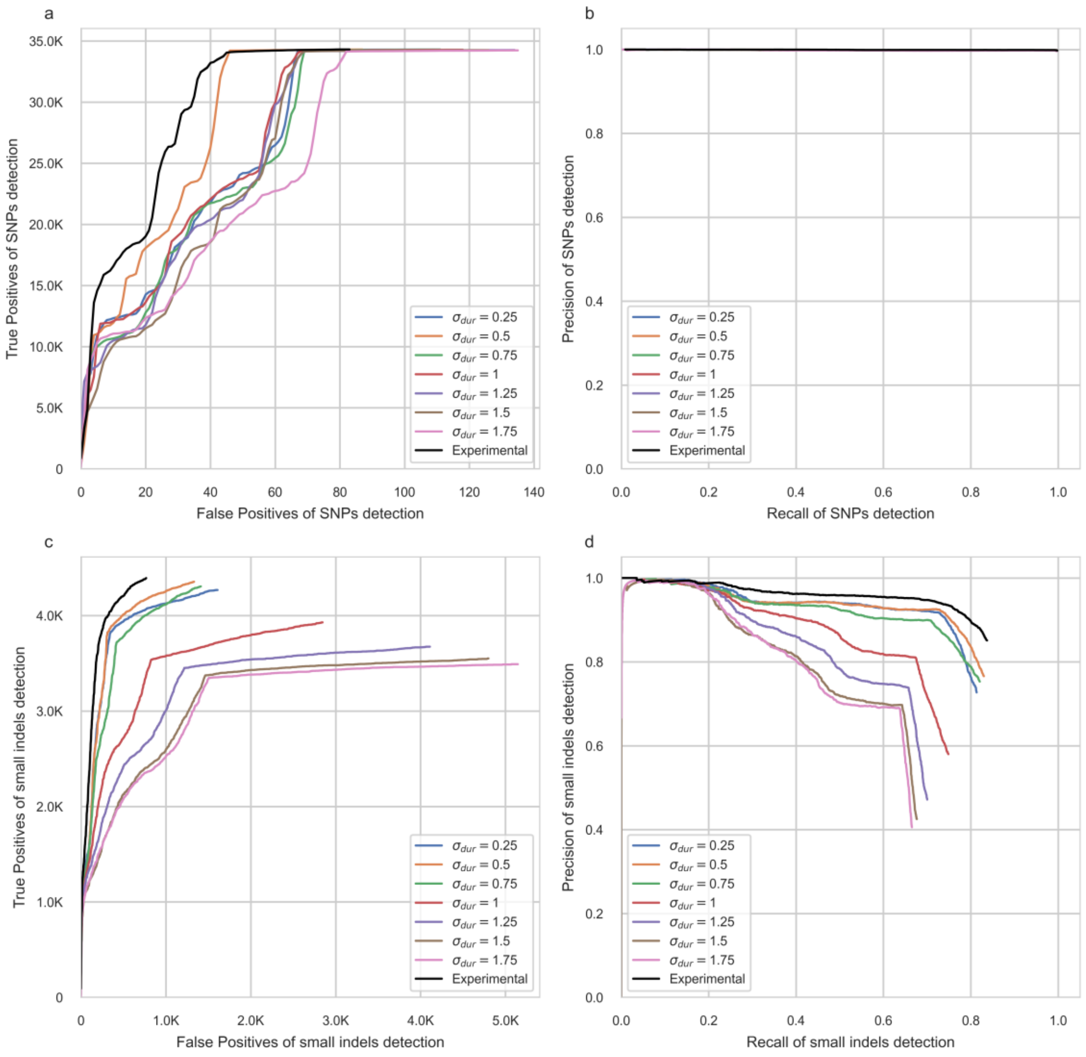

**Fig S2. Small variants detection performance for NanoSimFormer simulated signal using different event duration variance. a,** ROC curves for SNPs. **b,** Precision-recall curves for SNPs. **c,** ROC curves for small indels. **d.** Precision-recall curves for small indels.

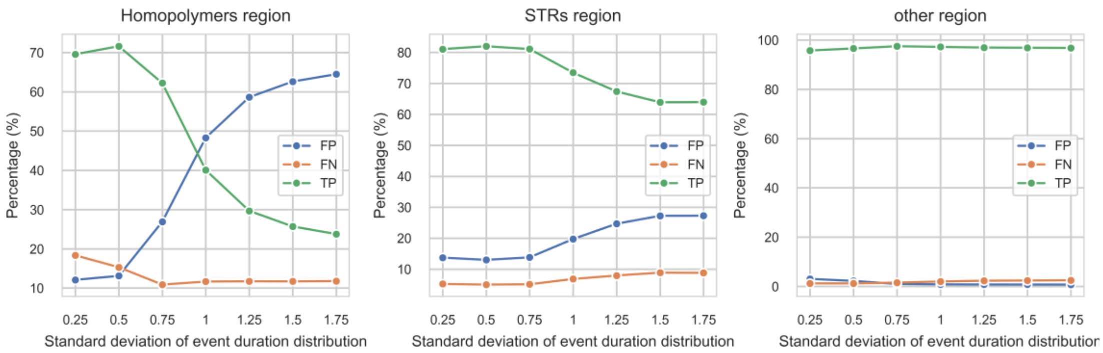

**Fig S3. Percentage of true positives (TP), false positives (FP), and false negatives (FN) of detected small variants in homopolymer, STRs and other regions of HG002 chr22 genome, for NanoSimFormer simulated signal using different event duration variance.**

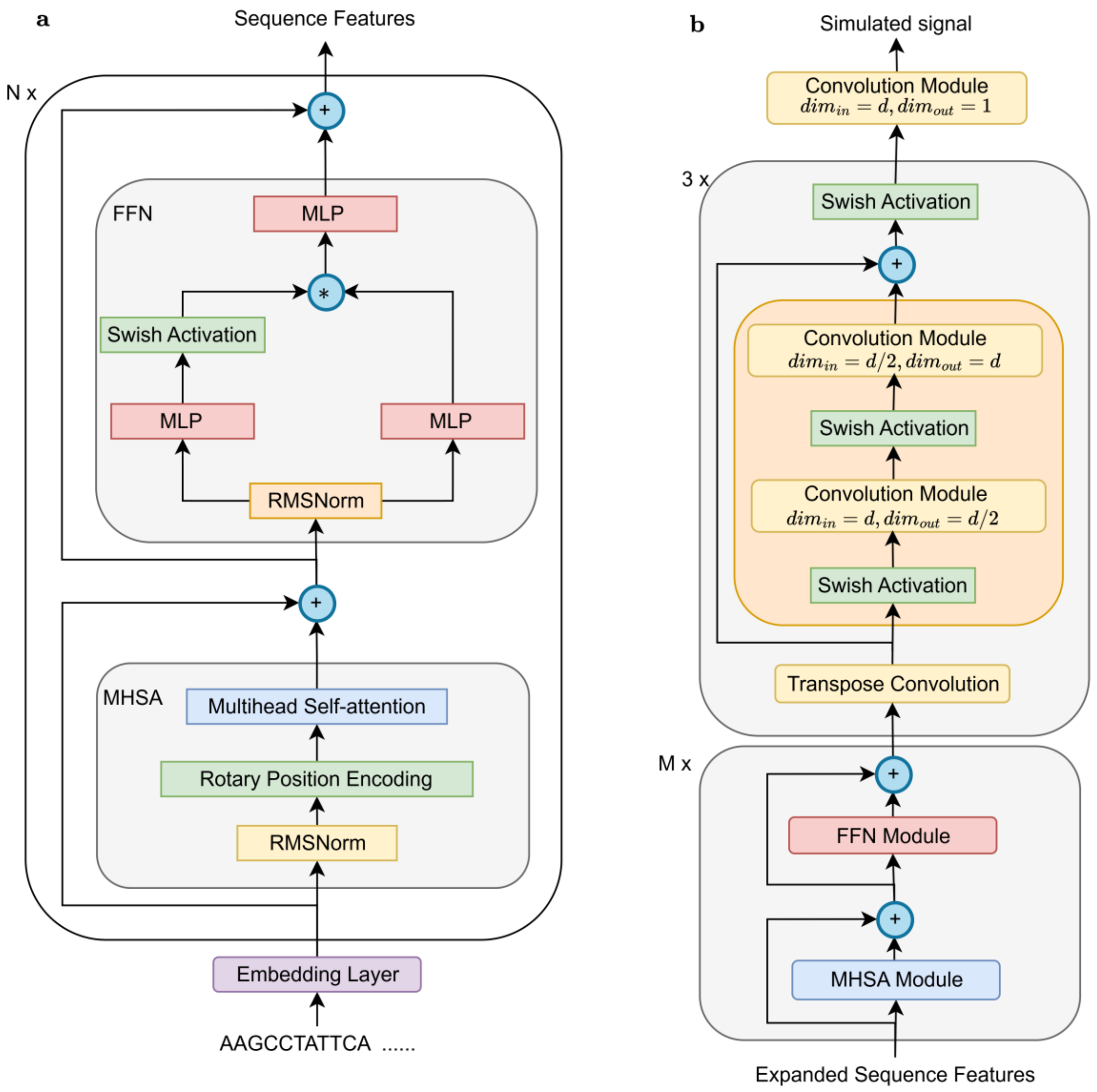

**Fig S4. Detailed architecture of sequence encoder and signal decoder in NanoSimFormer model.** **a,** Architecture of Transformer-based sequence encoder. **b,** Architecture of hybrid Transformer-CNN signal decoder.

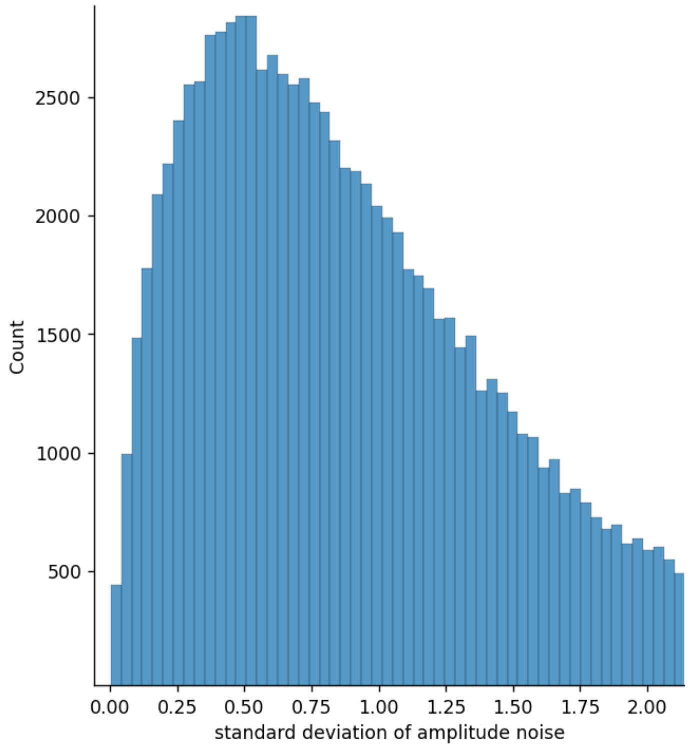

**Fig S5. Gamma distribution (shape=1.85, scale=0.55) of the amplitude noise standard deviation values used in the default simulation setting.**
